# Developmental olfactory dysfunction and abnormal odor memory in immune-challenged *Disc1^+/-^* mice

**DOI:** 10.1101/2024.05.17.594663

**Authors:** Fiona Parbst, Johanna K. Kostka, Anne Günther, Yu-Nan Chen, Ileana L. Hanganu-Opatz, Sebastian H. Bitzenhofer

## Abstract

Neuronal activity in the olfactory bulb (OB) drives coordinated activity in the hippocampal-prefrontal network during early development. Inhibiting OB output in neonatal mice disrupts functional development of the hippocampal formation as well as cognitive abilities. These impairments manifest early in life and resemble dysfunctions of the hippocampus and the prefrontal cortex that have been linked to neuropsychiatric disorders. Thus, we investigated a disease mouse model and asked whether activity in the OB might be altered, thereby contributing to the dysfunctional development of the hippocampal-prefrontal network. We addressed this question by combining in vivo electrophysiology with behavioral assessment of immune-challenged *Disc1^+/-^*mice that mimic the dual genetic-environmental etiology of neuropsychiatric disorders. In wildtype mice, we found high DISC1 expression levels in OB projection neurons during development. Furthermore, neuronal and network activity in the OB, as well as the drive from the bulb to the hippocampal-prefrontal network were reduced in immune-challenged *Disc1^+/-^* mice during early development. This early deficit did not affect odor-evoked activity and odor perception but resulted in impaired long-term odor memory. We propose that reduced endogenous activity in the developing OB contributes to altered maturation of the hippocampal-prefrontal network, leading to memory impairment in immune-challenged *Disc1^+/-^* mice.

## Introduction

Functional maturation of the central nervous system is shaped by neuronal activity during development [1–4]. This activity can be intrinsically generated within local networks or driven by input from the sensory periphery. For example, studies in rodents revealed that activity in the visual cortex during early development is critical for the refinement of the network and can be generated locally or driven by retinal waves that occur before eye opening [5,6]. Similarly, spontaneous activity from the cochlea is required for the maturation of the auditory cortex [7,8]. As for sensory areas, the maturation of higher association areas, such as the prefrontal cortex (PFC), is influenced by coordinated patterns of neuronal activity during development. Their perturbation during early postnatal development causes long-lasting circuit dysfunction, as result of an excitation-inhibition imbalance, and impairment of associated cognitive abilities [9,10].

However, what is driving early activity in the PFC is less clear. During development, inputs from the mediodorsal thalamus increase the activity level in the PFC and their inhibition impairs prefrontal maturation [11]. Monosynaptic projections from the intermediate and ventral hippocampus provide another excitatory drive for prefrontal activity during development [12,13]. A prominent candidate for the drive of developmental neuronal activity in these areas in turn is the olfactory system. From birth on, mouse pups rely on olfactory inputs to find the teats of the dam for feeding [14]. In line with this vital function, mitral and tufted cells, the projection neurons of the olfactory bulb (OB), as well as their axonal projections develop prenatally and their downstream connectivity is largely established at birth [15,16]. Consequently, coordinated neuronal activity in the OB emerges early in life and is more prominent in comparison to other brain areas during early postnatal development [17,18]. Even without a direct connection from the OB to the hippocampus or the PFC, strong projections from OB to the piriform cortex and lateral entorhinal cortex provide a short pathway by which the olfactory system can influence the hippocampal-prefrontal network [19–21]. In line with this, we previously showed that rhythmic activity in the OB entrains activity in the entorhinal cortex, hippocampus, and PFC already during early postnatal development [18,22]. Thus, activity in the olfactory system might play a similar role for the development of the hippocampal-prefrontal network as neuronal activity in the retina and the cochlea has for the maturation of the visual and the auditory system, respectively.

Indeed, we recently demonstrated that transient inhibition of OB outputs at the start of the second postnatal week in mice reduces coordinated activity in the hippocampal formation and impairs cognitive abilities later in life [23]. These findings resemble the developmental deficits observed in mouse models of neuropsychiatric disorders [24,25]. Accumulating evidence suggests an association of olfactory impairment with neuropsychiatric disorders. For example, inflammation of the olfactory epithelium, reduced OB volume, and deficits in odor perception have been reported for schizophrenia, psychosis, and depression [26–30]. However, it is unknown how alterations in the olfactory system might contribute to the pathogenesis of neuropsychiatric disorders that involve prefrontal-hippocampal dysfunction [24,31].

Here, we investigated the interactions of the olfactory system with the hippocampal-prefrontal network during neonatal development in a mouse model of neuropsychiatric disorders. Immune-challenged *Disc1^+/-^* mice are a dual-hit mouse model that combines two well-established models for neuropsychiatric disorders: a heterozygous mutation in the gene disrupted-in-schizophrenia 1 (*Disc1^+/−^*) resulting in a truncated DISC1 protein [32,33] and maternal immune activation by the viral RNA mimetic poly(I:C) [34]. This dual-hit gene-environment (GE) mouse model mimics the etiology of neuropsychiatric disorders and shows an impairment of coordinated activity in the hippocampal-prefrontal network during early development, as well as reduced performance in associated cognitive tasks [35–37].

## Results

### High developmental DISC1 expression in OB projection neurons

The DISC1 protein is involved in the regulation of several developmental processes, such as progenitor proliferation, neuronal migration, and synapse formation [38–40]. The regulation of the formation and maintenance of synaptic connections by DISC1 is considered particularly important in the context of its association with neuropsychiatric disorders [33,41]. In adult mice, DISC1 is expressed in several brain areas including the OB, the hippocampus, and the cerebral cortex [42–44].

As a first test for a potential role of the olfactory system in the pathophysiology of GE mice, we investigated DISC1 expression during postnatal development. Using immunohistochemistry, we found high expression levels of DISC1 in the OB of wildtype (WT) mice at the beginning of the second postnatal week (Fig 1A,B). DISC1 expression was significantly higher in the OB compared to the hippocampal subdivision CA1 (p=4.39×10^-4^) and the PFC (p=5.52×10^-4^) in WT mice. In GE mice, DISC1 expression was significantly reduced compared to WT controls in all areas (OB p=3.50×10^-4^, CA1 p=9.37×10^-5^, PFC p=1.7×10^-3^). More detailed examination of the OB revealed increased DISC1 expression in the neuronal processes but also in the cell bodies of neurons in the mitral cell layer (MCL) and the external plexiform layer (EPL) (Fig 1C).This is indicative for a strong DISC1 expression in mitral/tufted cells, the projections neurons of the OB. Immunostaining of OB slices of postnatal day (P) 10 WT mice, injected with the retrograde tracer CTB555 into the piriform cortex at P5, confirmed the expression of DISC1 in OB projection neurons (Fig 1D).

**Fig 1.**
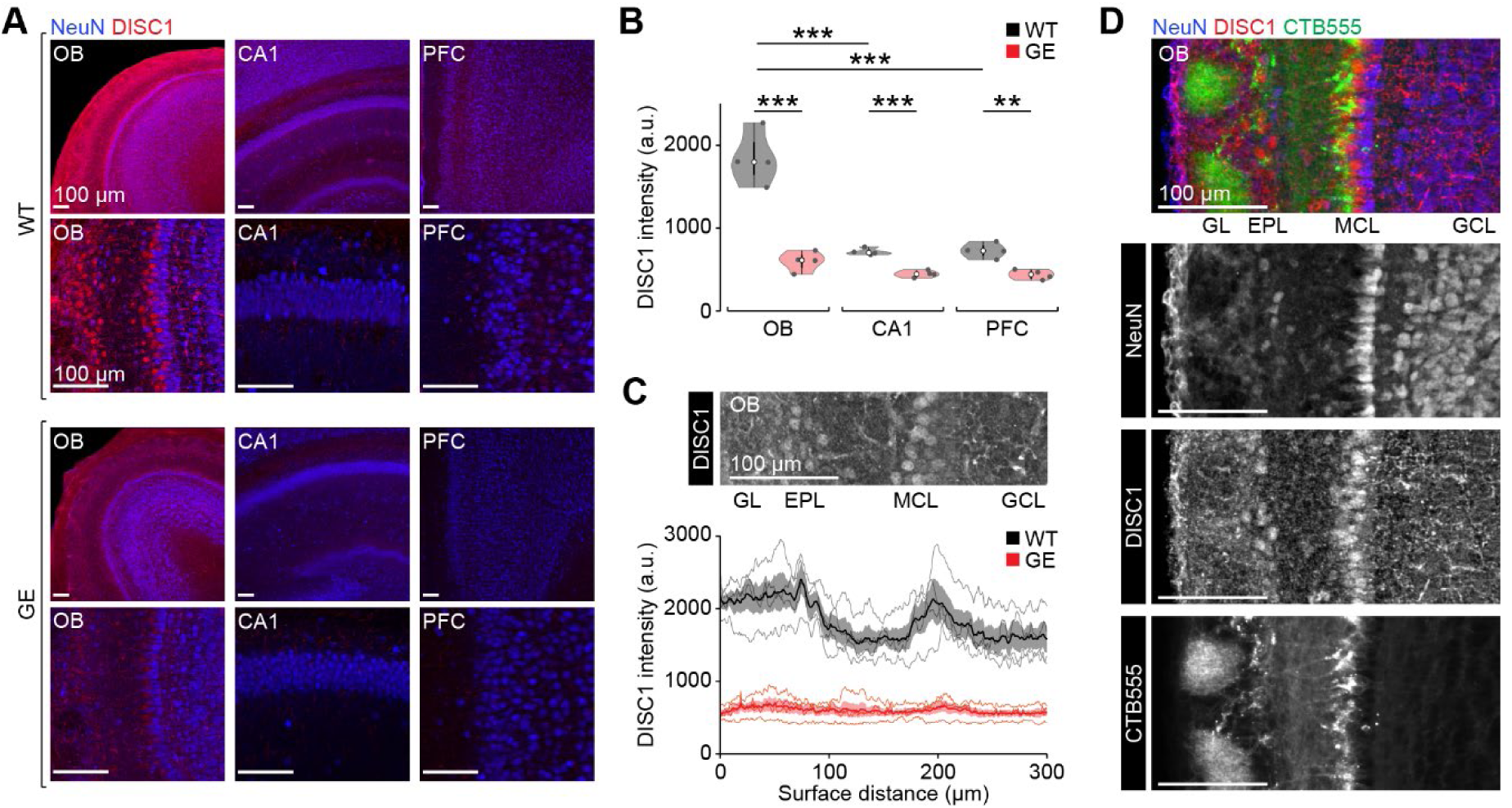
Strong expression of DISC1 in OB projection neurons at P10. **A** Coronal sections of OB, CA1, and PFC from P10 WT (top) and GE (bottom) mice immunostained for NeuN (blue) and DISC1 (red) at low and high magnification. Slices from different areas were stained in parallel and images were acquired with identical settings. **B** Fluorescence intensity of DISC1 immunolabeling in OB, CA1, and PFC in P9-10 WT (n=4) and GE (n=4) mice. **C** Top, coronal section of OB from a P10 WT mouse immunostained for DISC1. Bottom, spatially resolved DISC1 intensity in P9-10 WT (n=4) and GE (n=4) mice. **D** Coronal section of OB from a P10 WT mouse immunostained for DISC1 with OB projection neurons labeled by injection of the retrograde tracer CTB555 into the piriform cortex. Shaded areas in C correspond to standard error of the mean (SEM). Significant differences are indicated as *, **, *** for p<0.05, 0.01, 0.001, respectively. GL glomerular layer; GCL granule cell layer.

Together, these results support the idea of a potential developmental dysfunction of the OB in GE mice.

### Reduced OB activity in immune-challenged Disc1^+/-^ mice during development

To investigate developmental OB activity in GE mice, we performed *in vivo* electrophysiological recordings using multi-site silicon probes. We monitored endogenous and odor-evoked activity in the ventral OB of WT and GE mice at P8-10 (Fig 2A,B). The extracellular recordings were combined with respiration measurements using a pressure sensor. Both, WT and GE mice showed continuous activity in the local field potential (LFP) recorded in the OB with the typical dominant respiration rhythm (RR, 2-4 Hz) that reverses polarity at the MCL (Fig 2C). However, the power in RR and theta (4-12 Hz) frequency bands were significantly reduced in GE mice (RR p=5.1×10^-4^, theta p=0.004), whereas the power in beta (12-30 Hz) was not significantly different (p=0.09) (Fig 2D). Further, the firing rate of single units in the OB was reduced in GE mice, particularly for neurons in the MCL (MCL p=0.023; GCL p=0.51), indicating a reduction in the activity of OB projection neurons in the absence of odor stimulation (Fig 2E).

**Fig 2.**
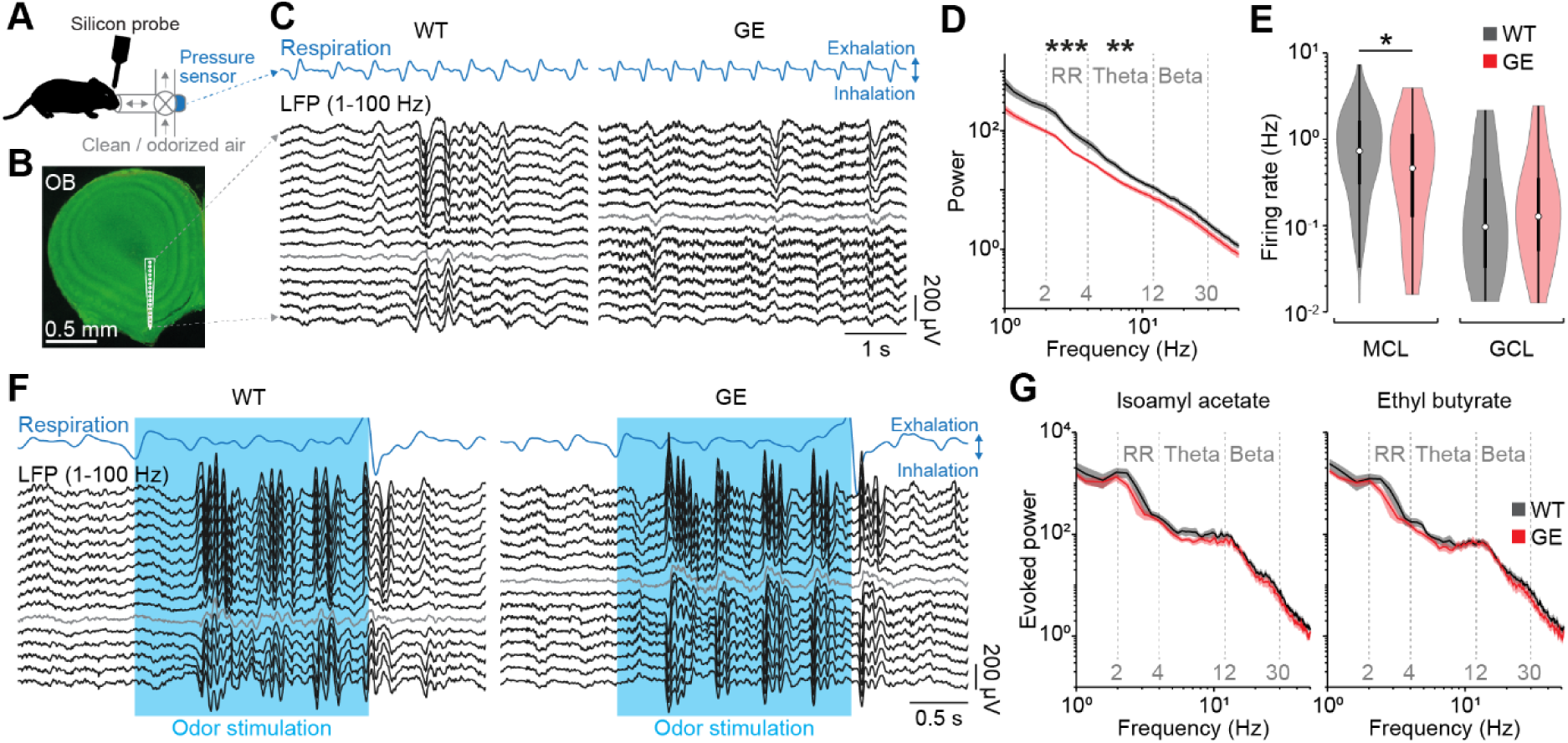
Reduced endogenous, but normal odor-evoked activity in the OB of immune-challenged *Disc1^+/-^* mice at P8-10. **A** Experimental setup for recordings of endogenous and respiration-triggered odor-evoked activity in the OB of P8-10 mice. **B** Example coronal section with a reconstruction of the DiI-labeled silicon probe tip in the ventral OB. **C** Example extracellular recording of endogenous activity from the ventral OB of a P10 WT and GE mouse using a silicon probe with 16 recording sites spanning across the MCL (gray). Down- and upward deflections on the respiration trace from the pressure sensor indicate inhalation and exhalation, respectively. **D** Power spectra of endogenous OB activity in P8-10 WT (n=17) and GE (n=14) mice. **E** Firing rate of endogenous OB activity in P8-10 WT and GE mice for units recorded in MCL (WT n=172, GE n=49 units) and GCL (WT n=55, GE n=38 units). **F** Same as C for odor-evoked activity. **G** Power spectra of odor-evoked OB activity in P8-10 WT (n=11) and GE (n=11) mice. Shaded areas in D,G correspond to SEM. Significant differences are indicated as *, **, *** for p<0.05, 0.01, 0.001, respectively.

Next, we used the respiration measurement for closed-loop respiration-triggered odor stimulation to investigate odor-evoked activity in the OB of WT and GE mice. Odor stimulation with the pure odorants isoamyl acetate or ethyl butyrate (1% v/v in mineral oil) was triggered by exhalations to guarantee for stable odor presentation at the subsequent inhalation. Odor presentation induced strong activation of the OB in WT and GE mice (Fig 2F). In contrast to endogenous activity, odor-evoked activity was similar for WT and GE mice across all frequency bands (Isoamyl acetate: RR p=0.088, theta p=0.36, beta p=0.32; Ethyl butyrate: RR p=0.12, theta p=0.29, beta p=0.39) (Fig 2G).

Notably, mice that only carry the genetic mutation (G) had a similar reduction to GE mice in the power of endogenous OB activity (G to WT: RR p=0.015, theta p=0.085, beta p=0.58; G to GE: RR p=0.75, theta p=0.82, beta p=0.69), whereas OB activity in mice that only received the environmental hit (E) of maternal immune activation was comparable to WT controls (E to WT: RR p=0.64, theta p=0.48, beta p=0.060; E to GE: RR p=0.056, theta p=0.82, beta p=0.26) (Fig 3).

**Fig 3.**
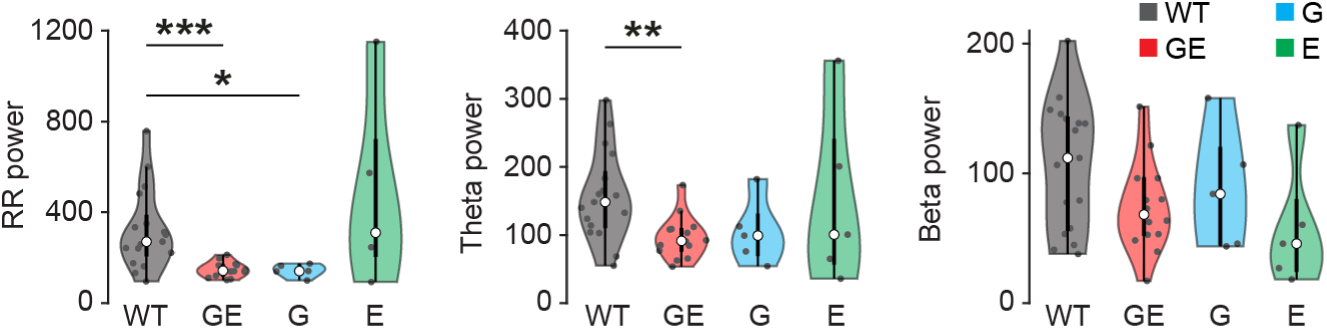
Reduced endogenous OB activity in *Disc1^+/-^*, but not in immune-challenged mice at P8-10. Power of endogenous OB activity quantified in different frequency bands (left, RR; middle, theta; right, beta) in P8-10 WT (n=17), GE (n=14), G (n=5), and E (n=5) mice. Significant differences are indicated as *, **, *** for p<0.05, 0.01, 0.001, respectively.

Thus, odor stimulation evokes activity in the OB of GE mice that is similar to WT controls whereas endogenous activity in the absence of odor stimulation is significantly reduced in GE mice.

### Reduced OB activity results in weaker drive of the hippocampal-prefrontal network

Does altered activity in the OB of GE mice affect activity in downstream areas? To address this question, we performed simultaneous recordings from OB, CA1 of the intermediate hippocampus, and the medial PFC of P8-10 WT and GE mice (Fig 4A,B). While OB activity was already continuous at this age, CA1 and PFC showed discontinuous patterns of electrical activity (Fig 4C) characteristic for this age [12]. As reported in our previous publications [35,36], prefrontal LFP power in theta and beta frequency range was reduced (theta p=8.6×10^-4^, beta p=0.0094) in GE mice when compared to WT controls, whereas no differences were detected for CA1 (theta p=0.19, beta p=0.49) (Fig 4D). Of note, the power in RR frequency band was reduced in both areas (CA1 p= 0.015, PFC p=0.0016) for GE mice in line with reduced RR power in the OB. Firing rates of single units in CA1 (p=0.41) and PFC (p=0.65) were similar for WT and GE (Fig 4E). In this study, mostly deep layers of the PFC were recorded, which explains the lack of altered firing rates that are characteristic for prefrontal layer 2/3 in neonatal GE mice [36].

**Fig 4.**
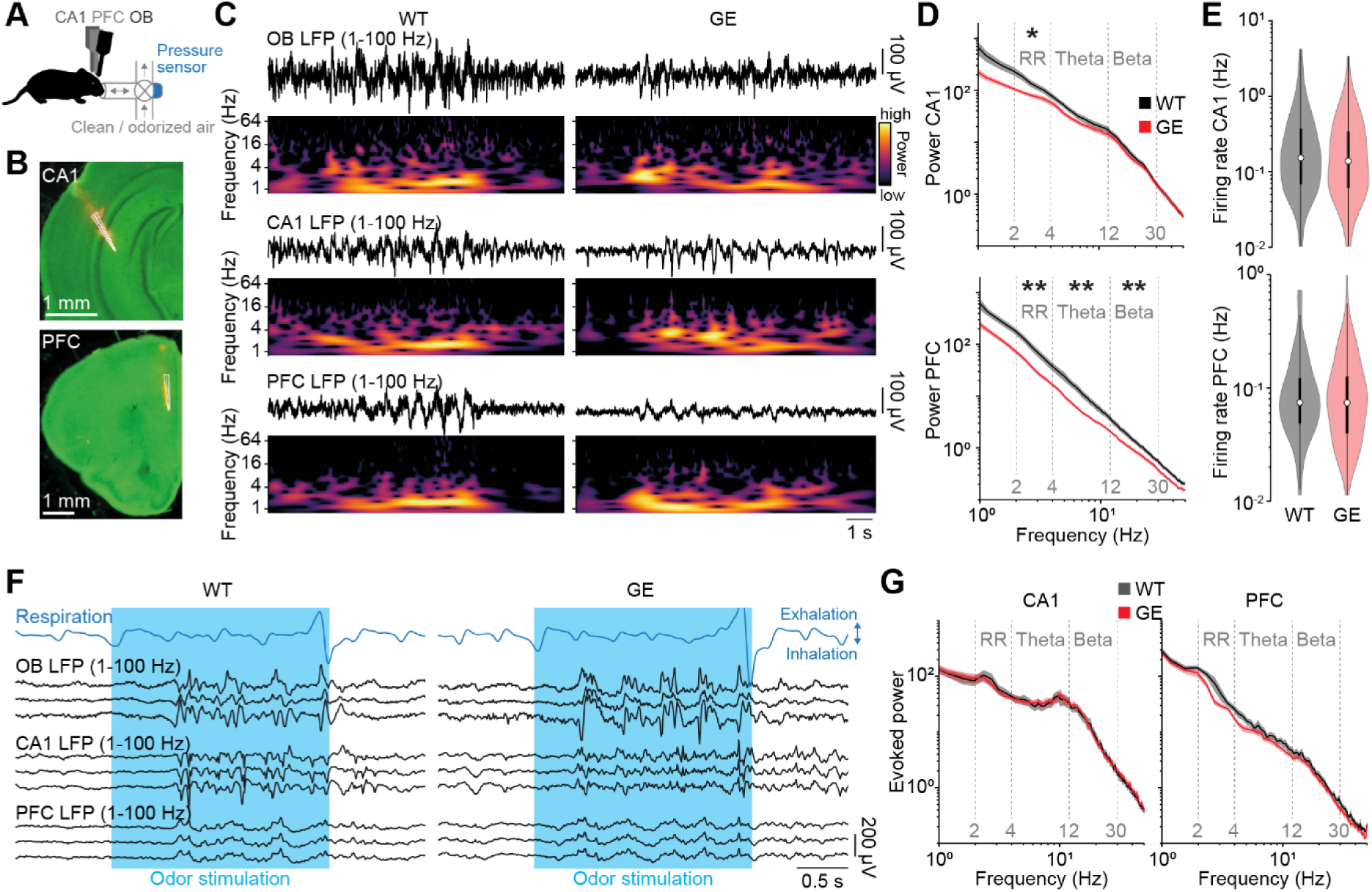
Reduced activity in the hippocampal-prefrontal network in immune-challenged *Disc1^+/-^* mice at P8-10. **A** Experimental setup for triple recordings of endogenous and respiration-triggered odor-evoked activity in OB, CA1, and PFC of P8-10 mice. **B** Example coronal sections with a reconstruction of the DiI-labelled silicon probe tips in CA1 of the intermediate hippocampus (top) and the medial part of the PFC (bottom). **C** Examples of endogenous LFP activity recorded simultaneously from OB, CA1, and PFC of a P10 WT and GE mouse and the corresponding wavelet spectra. **D** Power spectra of endogenous CA1 and PFC activity in P8-10 WT (CA1 n=15, PFC n=14) and GE (CA1 n=14, PFC n=14) mice. **E** Firing rate of endogenous CA1 and PFC single unit activity in P8-10 WT (CA1 n=270, PFC n=167 units) and GE (CA1 n=182, PFC n=153 units) mice. **F** Examples of odor-evoked LFP activity recorded simultaneously from OB, CA1, and PFC of a P10 WT and GE mouse. **G** Power spectra of odor-evoked CA1 and PFC activity in P8-10 WT (n=11) and GE (n=11) mice. Shaded areas in D,G correspond to SEM. Significant differences are indicated as *, **, *** for p<0.05, 0.01, 0.001, respectively.

Next, we tested whether there is a difference in the propagation of odor-evoked activity to the hippocampal-prefrontal network in GE mice. Odor stimulation evoked pronounced activation in CA1 and PFC of WT and GE mice (Fig 4F). As for OB, we found no significant differences for odor-evoked activity in CA1 (RR p=0.92, theta p=0.86, beta p= 0.75) and PFC (RR p=0.062, theta p=0.28, beta p= 0.55) of P8-10 WT and GE mice (Fig 4G).

Thus, similar to the OB, endogenous activity in CA1 and PFC of P8-10 GE mice is decreased, yet odor-induced activity is similar to age-matched WT controls.

Reduced RR power in OB, CA1, and PFC of GE mice during development suggests a possible reduction in the drive from the OB to the hippocampal-prefrontal network. To test this hypothesis, we used pairwise analyses of the simultaneously recorded areas to test their functional interactions in P8-10 WT and GE mice. First, we calculated the amplitude correlation of the LFP between areas as a measure for non-directed interactions that is based on the similarity of the amplitude fluctuations at a given frequency. We found a significant reduction in the interaction between OB and CA1 (RR p=0.018, theta p=0.012, beta p=0.49), OB and PFC (RR p=0.045, theta p=0.023, beta p=0.063), as well as CA1 and PFC (RR p=0.011, theta p=0.039, beta p=0.37) in RR and theta frequency but not in beta frequency for GE mice (Fig 5A). Next, we tested more directed measures of functional interactions.

**Fig 5.**
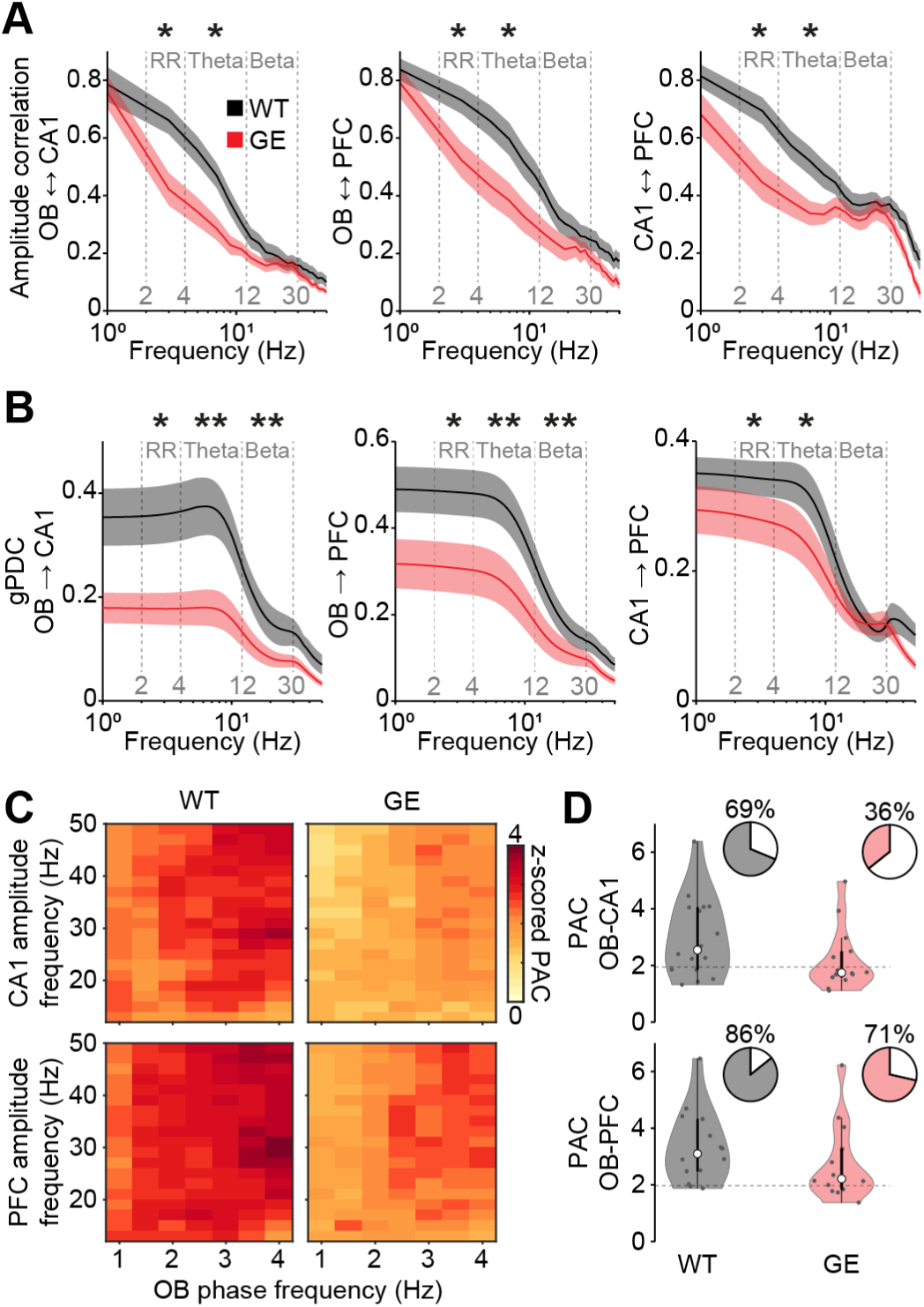
Reduced drive of the hippocampal-prefrontal network in immune-challenged *Disc1^+/-^* mice at P8-10. **A** Frequency-resolved amplitude correlation between OB-CA1 (WT n=16, GE n=14), OB-PFC (WT n=14, GE n=14), and CA1-PFC (WT n=13, GE n=14) for P8-10 WT and GE mice. **B** Frequency-resolved gPDC from OB to CA1 (WT n=16, GE n=14), from OB to PFC (WT n=14, GE n=14), and from CA1 to PFC (WT n=13, GE n=14) for P8-10 WT and GE mice. **C** Color-coded average PAC of CA1 (WT n=16, GE n=14) and PFC (WT n=14, GE n=14) LFP amplitude at fast frequencies (12-50 Hz) to slow frequency oscillations (1-4 Hz) in OB for P8-10 WT and GE mice. **D** Z-scored PAC of CA1 and PFC LFP amplitude at fast frequencies to slow frequencies in the OB for P8-10 WT and GE mice. Pie charts show the percentage of recordings with significant coupling. Dotted lines correspond to a z-score of 1.96 indicating the significance level. Shaded areas in A,B correspond to SEM. Significant differences are indicated as *, **, *** for p<0.05, 0.01, 0.001, respectively.

Generalized partial directed coherence (gPDC), which assesses the directionality of pairwise interactions, revealed a strong reduction in the drive from OB to CA1 (RR p=0.011, theta p=0.0044 beta p=0.0037) and OB to PFC (RR p=0.011, theta p=0.0051 beta p=0.0016) in all frequency bands for GE mice (Fig 5B). The directed interaction from CA1 to PFC was reduced in RR (p=0.024) and theta (p=0.018) but not beta (p=0.76) frequency for GE mice, as previously reported [37]. Finally, we quantified phase-amplitude coupling (PAC) to evaluate the cross-frequency modulation of oscillatory power at fast frequencies (12-50 Hz) in CA1 and PFC by the slow RR generated in the OB. The strength of PAC from OB to CA1 and PFC was reduced in GE mice, as was the percentage of recordings with significant cross-frequency coupling (OB to CA1: WT 11 of 16, GE 5 of 14 mice; OB to PFC: WT 12 of 14, GE 10 of 14 mice) (Fig 5C,D).

Together, these findings show that the drive from OB to CA1 and PFC is reduced in GE mice.

### Normal odor detection, but impaired odor memory in immune-challenged Disc1^+/-^ mice

Normal propagation of odor-evoked activity from OB to the hippocampal-prefrontal network suggests that GE mice might have normal odor processing during development. To address this hypothesis, we recorded ultrasonic vocalizations (USV) of P9 WT and GE mice when exposed to the odorant citral. Citral triggers an innate aversive response and reduces USV calls in neonatal mice, similar to the odor of adult males [45]. Pups were placed in a small chamber with a continuous flow of clean air for a baseline period after which citral was added to the airstream (Fig 6A-C). This procedure was repeated with increasing concentrations of citral. While concentrations of 0.0001% (WT p=0.50, GE p=0.17) and 0.01% (WT p=0.37, GE p=0.40) of citral (v/v in mineral oil) did not reduce call rates, both, WT and GE mice similarly reduced their call rate in response to citral at 1% (WT p=1.1×10^-4^, GE p=3.4×10^-6^, WT vs. GE p=0.32) (Fig 6D).

**Fig 6.**
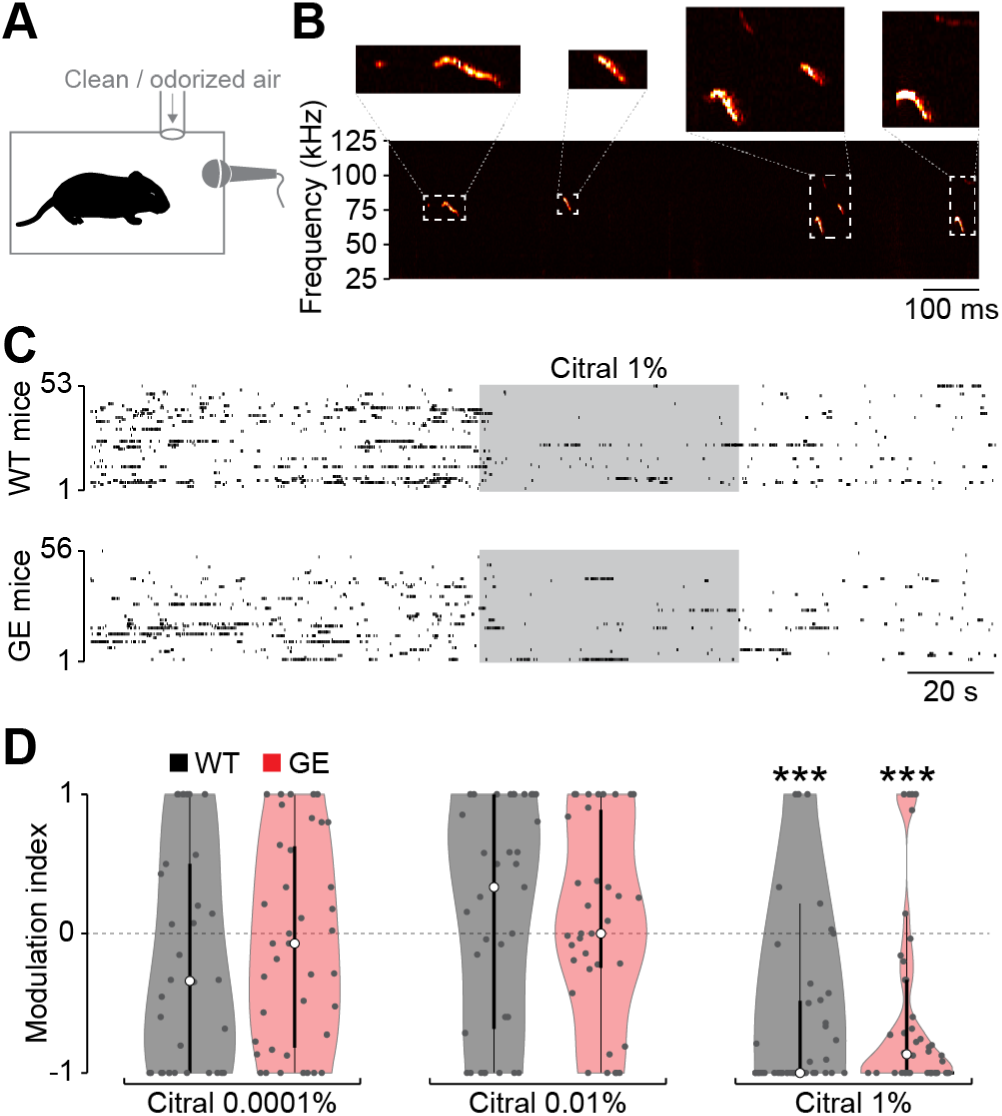
Odor detection is intact in immune-challenged *Disc1^+/-^* mice at P9. **A** Experimental setup for USV recordings during odor exposure. **B** Example spectrogram of USVs of a P9 mouse. **C** Raster plot of USV suppression in response to the odorant citral for P9 WT (n=53) and GE (n=56) mice. Each line represents one mouse. **D** Modulation index of USV numbers in response to the odorant citral at different concentrations. Significant differences are indicated as *, **, *** for p<0.05, 0.01, 0.001, respectively.

Thus, simple odor detection is not impaired in developing GE mice, consistent with normal odor-evoked activity.

We recently reported that transient inhibition of OB outputs from P8 to P10 in WT mice perturbs the functional maturation of the hippocampal formation and results in long-lasting cognitive deficits [23]. Thus, we hypothesized that the reduced endogenous OB activity and drive to the hippocampal-prefrontal network in developing GE mice might cause similar impairments. We used neonatal odor learning, a one-trial associative odor learning task for mouse pups [46], to test the learning and memory abilities in P10-11 WT and GE mice (Fig 7A). For this test, the dam was separated from the pups for 2 hours. Subsequently, a novel odor was applied to the teats before the dam was returned to the pups. The separation period guarantees feeding of the pups soon after odor application such that an association between food consumption and the odor can be formed. After a second separation period, pups were tested in an odor-place preference test with the learned test odor and a novel control odor (Fig 7B). Isoamyl acetate and ethyl butyrate (1% in mineral oil) were randomly assigned as test and control odor for each litter and the position of the test odor was randomized for each pup.

**Fig 7.**
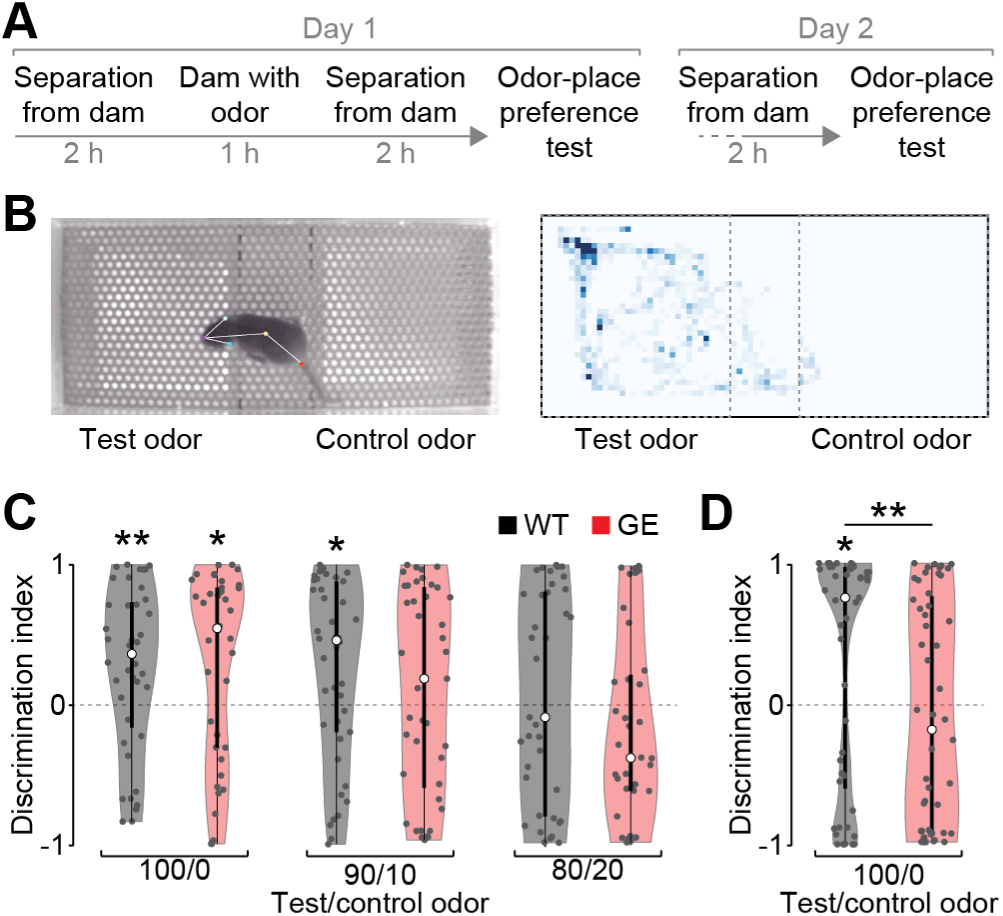
Impaired long-term odor-memory in immune-challenged *Disc1^+/-^* mice at P10/11. **A** Timeline of neonatal odor learning and odor-place preference test for P10-11 mice. **B** Left, example image of a P10 mouse in the odor-place preference test chamber. Right, example color-coded position of a mouse’s nose traced by DeepLabCut during the odor-place preference test. **C** Discrimination index of test and control odor in the odor-place preference test on the same day of neonatal odor learning for P10 WT and GE mice. Test and control odors were presented pure (100/0, WT n=40, GE n=38 mice) or mixed (90/10, WT n=42, GE n=41 mice; 80/20, WT n=37, GE n=36 mice). **D** Discrimination index of test and control odor in the odor-place preference test the day after neonatal odor learning for P11 WT (n=57) and GE (n=52) mice. Significant differences are indicated as *, **, *** for p<0.05, 0.01, 0.001, respectively.

Both, WT and GE mice spend more time on the side of the test odor when pure odors were presented in the odor-place preference test (WT p=0.0075, GE p=0.021) and no difference was found between the groups (p=0.73) (Fig 7C). To increase the difficulty of the test, mice were presented with a mixture of test and control odors during the odor-place preference test. Interestingly, only WT controls were able to distinguish the learned test odor at a mixture of 90/10% (WT p=0.015, GE p=0.31) but there was no significant group difference for WT and GE mice (p=0.39). Neither WT nor GE mice showed a place preference when odors were presented at a mixture of 80/20% (WT p=0.48, GE p=0.22). These results indicate that odor-learning is not impaired in GE mice at P10. However, when the odor-place preference test was done with a 24h delay period after neonatal odor learning, WT pups strongly preferred the learned test odor (p=0.012), whereas GE mice did not distinguish between test and control odor (p=0.18) and their performance was significantly reduced compared to WT mice (p=0.0026) (Fig 7D).

Together, these data revealed that odor detection and learning are largely normal in developing GE mice, whereas their long-term memory is impaired.

## Discussion

In this study, we examined early OB activity and its influence on developing hippocampal-prefrontal networks in GE mice. We found strong DISC1 expression in OB projection neurons during development, which was significantly reduced in GE mice. GE pups displayed reduced endogenous activity in the OB, whereas odor-evoked activity was comparable to WT controls. Correspondingly, the drive from the OB to the hippocampal-prefrontal network was reduced for endogenous activity, but the propagation of odor-evoked activity to HP and PFC was not altered. Consistent with these findings, odor detection and learning were not affected in GE mouse pups, but their long-term odor memory was impaired. We conclude that reduced endogenous activity in the OB contributes to the altered maturation of the hippocampal-prefrontal network as well as memory deficits in GE mice.

During development, endogenous and sensory-evoked activity is required for the refinement of immature neuronal networks. Neuronal activity influences a range of developmental processes such as neuronal survival and dendritic growth, as well as synapse formation and pruning [1–3,10]. The endogenous and odor-induced OB activity is a major drive for developing hippocampal-prefrontal networks [18,22,47]. Transient inhibition of OB outputs at the beginning of the second postnatal week disrupts the functional maturation in the hippocampal formation, which causes long-lasting deficits in cognitive abilities [23]. Here, we show that the reduction of endogenous activity in the OB of GE mice appears to have similar consequences for the maturation of the hippocampal-prefrontal network. We found a significant reduction in the activity of OB projection neurons and a concurrent decrease in the coordinated activity in slow oscillatory rhythms in GE mice. Particularly, the power in RR, oscillatory activity within 2-4 Hz, was significantly reduced in OB, as well as in CA1 and PFC. This rhythm is driven by repetitive input to the OB generated by the inhalation-exhalation cycle and coordinates activity in downstream areas in neonatal and adult mice [18,22,48–50]. Reduced coordination of activity in CA1 and PFC as a result of a reduced coordination of OB activity by respiration in GE mice might underlie the detrimental effect on the maturation of the hippocampal-prefrontal network. Interestingly, the single-hit mouse model carrying the genetic disruption of *Disc1^+/-^* without the environmental hit showed a similar reduction of RR power in OB, indicating that the genetic deficit alone is sufficient for RR impairment. In contrast, oscillatory activity in CA1 and PFC is only affected when the *Disc1^+/-^*mutation is combined with maternal immune activation [51]. This distinction might be linked to the strong expression of DISC1 in the developing OB of WT mice.

DISC1 is also expressed in CA1 and PFC during development and previous studies found that specific knockdown of DISC1 in these areas in combination with maternal immune activation suffices to impair hippocampal-prefrontal activity [52,53]. However, strong DISC1 expression in OB projection neurons and high levels of OB activity during development in WT mice indicate a particular role of reduced OB activity in disturbed maturation of the hippocampal-prefrontal network in GE mice. The clear directionality in the drive of activity from OB to CA1 and PFC is consistent with anatomical data showing that feedback projections to the OB develop late and are still sparse at the beginning of the second postnatal week (Kostka & Bitzenhofer, 2022).

The axons of OB projection neurons are bundled in the lateral olfactory tract that distributes olfactory information to a range of brain areas [19]. While the OB has no direct projections to CA1 and PFC in mice, strong projections through the piriform cortex and the lateral entorhinal cortex provide a short pathway from the OB to the hippocampal-prefrontal network. This pathway is functional early during development [18,22,47] and, notably, impaired activity in the lateral entorhinal cortex has been found in GE mice [37]. In adult mice, inhibition of the lateral entorhinal cortex impairs performance in odor discrimination tasks [54] and odor-context learning [55]. This is not to say that odor detection or discrimination happens in the lateral entorhinal cortex but it shows that this pathway from OB to the hippocampal-prefrontal network is critical for the execution of odor-related tasks. Parallel pathways, such as direct projections from the anterior olfactory nucleus and the lateral entorhinal cortex to the PFC [37,56], might provide alternative routes for olfactory information to higher associative areas. However, the pathway through the lateral entorhinal cortex might be particularly vulnerable to reduced OB activity during early postnatal development as indicated by lasting morphological and functional alterations of entorhinal neurons after transient inhibition of OB projection neurons [23], as well as in GE mice [57].

For now, we can only speculate how these findings relate to deficits of the olfactory system reported for neuropsychiatric disorders. Olfactory impairment has been suggested as an early indicator for several neuropsychiatric disorders, such as schizophrenia and psychosis [26–28]. Patients with first episode psychosis were shown to display deficits in odor identification tasks, and a reduced OB volume and inflammation of the olfactory epithelium have been associated with schizophrenia [28,30]. We found largely normal odor detection and odor learning in GE mice consistent with normal odor-evoked activity during development, but long-term odor memory was impaired. More rigorous behavioral testing might reveal subtle olfactory dysfunctions early on but the options for behavioral tests in neonatal mice are limited. Alternatively, altered functional maturation of the OB and the feedback projections from higher association areas might accumulate throughout development and only result in olfactory deficits later in life. Future investigations deepening the present approaches could provide valuable insights on how olfactory deficits might act as valuable early diagnostic markers in neuropsychiatric disorders.

## Materials and methods

### Ethical approval

All experiments were performed in compliance with German laws and guidelines of the European Union for the use of animals in research (EU Directive 2010/63/EU) and were approved by the local ethical committee (Behörde für Gesundheit und Verbraucherschutz Hamburg, G17/015, N18/015).

### Animals

Timed-pregnant mice from the University Medical Center Hamburg-Eppendorf animal facility were housed individually in a 12/12 h light/dark cycle and had access to water and food ad libitum. The day of vaginal plug detection was defined as gestational day 0.5, and the day of birth was defined as postnatal day (P)0. Experiments were performed on pups of both sexes during neonatal development (i.e. P8–P11).

Pregnant *Disc1* mice (B6.129S6-Disc1^tm1Kara^) carrying a mutation resulting in a truncated transcript on a C57BL6/J background [32] received viral RNA mimetic poly(I:C) (25 mg/kg) injected intraperitoneally (i.p.) at gestational day 9.5 to induce maternal immune activation. Immune-challenged *Disc1^+/−^* mice (referred to as GE) combine genetic and environmental risk factors in the pathogenesis of neuropsychiatric disorders. Offspring of C57BL/6J mice were used as wild-type control animals (referred to as WT).

### Histology

P9-10 mice were anesthetized with 10% ketamine (aniMedica, Germany) / 2% xylazine (WDT, Germany) in 0.9% NaCl (10 μg/g body weight, intraperitoneal) and transcardially perfused with 4% paraformaldehyde (Histofix, Carl Roth, Germany). Brains were removed and postfixed in 4% paraformaldehyde for 24 h. Brains were sectioned coronally with a vibratome at 100 μm for immunostaining.

#### Immunostaining

Free-floating slices were permeabilized and unspecific binding sites were blocked in PBS containing 0.8% Triton X-100 (Sigma-Aldrich, MO, USA), 5% normal goat serum, and 5% normal donkey serum (Jackson Immuno Research, PA, USA) for 1 h. Slices were incubated with primary antibodies in 0.8% Triton X-100, 1% normal goat serum, and 1% normal donkey serum for 3d at 4°C. Subsequently, slices were washed in PBS and incubated with secondary antibodies for 3 h at room temperature. Slices were washed and transferred to glass slides, before being covered with Vectashield(Vectorlabs, CA, USA). Images of immunostainings were acquired with a confocal microscope (FV1000, Olympus, Japan). Images were processed and analyzed with ImageJ.

#### Retrograde tracing

P5 mice were injected with the retrograde tracer CTB555 (200 nl at 100 nl/min, cholera toxin subunit B, Alexa Fluor 455 conjugate) into the piriform cortex under isoflurane anesthesia (induction: 5%, maintenance: 2%). Pups were transcardially perfused and brains were removed for immunostaining at P10.

### Surgical procedure for electrophysiology

For *in vivo* electrophysiological recordings, P8-10 mice underwent surgery under isoflurane anesthesia (induction: 5%, maintenance: 2%). The skin above the skull was removed and local anesthetic (0.5% bupivacaine/1% lidocaine) was applied on the neck muscles. Two plastic bars were fixed on the nasal and occipital bones with dental cement. Craniotomies of about 0.5 mm diameter were performed above the right OB (0.5-0.8 mm anterior to the frontonasal suture, 0.5 mm lateral to the internasal suture), the CA1 subdivision of the intermediate hippocampus (2.5 mm anterior to lambda, 3.5 mm lateral to the midline), and the medial part of the PFC (0.5 mm anterior to bregma, 0.1-0.5 mm lateral to the midline). Throughout surgery, recovery, and recording mice were kept on a heating blanket at 37°C.

### Electrophysiological recordings

Extracellular recordings were performed simultaneously from the ventral OB, hippocampal CA1, and PFC in non-anesthetized P8-10 mice. For this, one-shank silicon probes (NeuroNexus, MI, USA) with 16 recording sites (50 µm inter-site spacing) were inserted into OB (0.5-1.8 mm deep, angle 0°), CA1 (1.3-1.9 mm deep, angle 20°), and PFC (1.8-2.1 mm deep, angle 0°). Before insertion, the electrodes were covered with DiI (1,1’-Dioctadecyl-3,3,3’,3’-tetramethylindocarbocyanine perchlorate, Molecular Probes, Eugene, OR) for confirmation of electrode position post mortem. A silver wire was inserted into the cerebellum and served as ground and reference electrode. A recovery period of 20 min after the insertion of electrodes was provided before data acquisition. Extracellular signals were band-pass filtered (0.1–9000 Hz) and digitized (32 kHz) with a multichannel extracellular amplifier (Digital Lynx SX; Neuralynx, Bozeman, MO, USA) and the Cheetah acquisition software (Neuralynx). After recordings, mice were deeply anesthetized with 10% ketamine/2% xylazine in 0.9% NaCl solution (10 µg/g body weight, i.p.) and transcardially perfused with Histofix (Carl Roth, Karlsruhe, Germany) containing 4% paraformaldehyde for subsequent identification of electrode positions in coronal slices.

### Olfactory stimulation

A custom-made, Arduino-controlled olfactometer with a constant stream of clean air (0.9 L/min) to the nose was used to present odors to the animals. Odors were presented for 2 s triggered by the respiration cycle of mice to ensure constant odor concentration at the first odor inhalation. Two different odors (ethyl butyrate and isoamyl acetate, 1% in mineral oil) were delivered in a randomized order for 40 repetitions each.

### Analysis of electrophysiological data

Electrophysiological data were analyzed with custom-written algorithms in Matlab R2021a environment. For LFP analysis data were band-passfiltered (1-100 Hz) using a phase preserving third-order Butterworth filter. For LFP data recorded in the OB, the recording site centered in the EPL was used, whereas, for the analysis of spiking activity, recording sites in the MCL or GCL were considered. For the analysis of hippocampal LFP, a recording site located in CA1 below the pyramidal layer was selected, while for the analysis of spiking activity, all recording sites located in CA1 were used. For LFP analysis in the PFC, a recording site centered in the prelimbic region was considered and spiking activity from all recording sites was included.

#### Power spectral density

Power spectral density was calculated using Welch’s method with non-overlapping windows of 2 s for endogenous activity or for a 2 s window for odor stimulation. Time-frequency power plots were calculated with a continuous wavelet transform (Morlet wavelet). Frequency bands for statistical comparisons were defined as RR (2-4 Hz), theta (4-12 Hz) and beta (12-30 Hz).

#### Frequency resolved amplitude correlation

LFP from OB, CA1 and PFC was band-pass filtered in frequency bins of 2 Hz from 1 to 30 Hz and Hilbert transformed to extract the absolute amplitude. Subsequently, pairwise Pearson correlation coefficients of frequency resolved envelopes were calculated for OB, CA1, and PFC.

#### Generalized partial directed coherence

gPDC was calculated in the frequency domain to investigate the directional interaction between areas. This linear Granger causality measure is based on the decomposition of multivariate partial coherence computed from multivariate autoregressive models. LFP signals of 1 s length were denoised using the MATLAB wavelet toolbox (ddencmp.m and wdencmp.m) before gPDC was calculated with a previously described algorithm [58].

#### Phase–amplitude coupling

Cross-frequency coupling was calculated between the phase of the slow frequency in OB and the amplitude at fast frequencies (12-80 Hz) in CA1 and PFC according to a previously described algorithm [59]. Band-pass filtered LFP was Hilbert transformed to extract the phase and amplitude. The amplitude of the 12 to 50 Hz filtered LFP in CA1 and PFC was determined at each phase of the filtered OB signal. PAC matrices were z-scored and the average was calculated for RR (2–4 Hz) phase to higher frequencies (13-50 Hz) coupling.

#### Spiking analysis

Single units were automatically detected and clustered using the python-based software klusta [60] and manually curated using phy (https://github.com/cortex-lab/phy). For OB recordings, units detected around the channel (+/1 1 channels) where the RR reverses in polarity were considered for MCL spiking activity, whereas units in channels >3 channels central from the MCL were considered for GCL spiking activity.

### Behavior

#### Neonatal odor detection

The suppression of ultrasonic vocalizations (USVs) of neonatal mice when exposed to the odor citral was used to test for neonatal odor detection. For each test, a P9 mouse was removed from the home cage, placed in a soundproof test box, and allowed to accommodate for 120 s. Pressurized air was pumped through the box at a rate of 2 L/min. USVs were recorded with an ultrasonic microphone (Avisoft UltraSoundGate, Avisoft Bioacoustics) at a sampling rate of 250 kHz for 90 s with clean air, 60 s with citral, and 60 s of clean air, followed by a 60 s break. Each test consisted of three consecutive trials with increasing concentration of citral at 10^-4^, 10^-2^, and 1% diluted in mineral oil. USVs from 25 to 125 kHz were detected using DeepSqueak [61].

#### Neonatal Odor Learning

Neonatal odor learning of P10 mice was assessed with a modified version of a previously established protocol [46]. For one-trial associative odor learning, the dam was removed from the home cage for 2 h before the test odor was applied to the teats of the dam with a saturated cotton swab and she was placed back to the home cage for 1 h. Isoamyl acetate and ethyl butyrate (1% mineral oil) were used randomly as test and control odors per litter. The dam was removed again for 2 h before the pups were tested in an odor place preference test. The test arena consisted of a rectangular acrylic chamber (17.5 × 6.5 × 6.5 cm) with metal grid flooring, divided into two 6.5 cm odor zones at the ends and a 4.5 cm neutral zone in the center. Odor zones were odorized by placing acrylic trays beneath the grid flooring with 500 µl of either the test odor or a control odor on a filter paper. Test and control odors were randomized between the two odor zones. For the test, a pup was placed in the center of the arena (i.e., neutral zone) and video-taped from above for 3 min using a camera (UI 2250-SE-M, IDS GMBH). Between each test, the chamber was cleaned with ethanol and allowed to dry. Odor place preference tests were performed with pure odors (1% in mineral oil) or odor mixtures (90/10, 80/20 test/control odor). Mice were tracked using DeepLabCut [62]. The time spent over the different zones was quantified and analyzed in Matlab.

To assess long-term neonatal odor memory, the dam was removed for 2 h on the following day (P11) and the odor place preference test was repeated with pure test versus control odor (1% in mineral oil).

### Statistics

Statistical analysis was performed in Matlab R2021a environment. Data were tested for normal distribution. Paired and unpaired t-tests were used for normally distributed data, whereas non-parametric Wilcoxon rank sum and sign rank tests were used for non-normally distributed data to test for significant differences. Data are presented as violin plots or as mean ± SEM. Significance levels of P < 0.05 (*), P < 0.01 (**) or P < 0.001 (***) were considered.

## Acknowledgements

We thank Dr. Joseph Gogos for providing the Disc1 mice. We thank A. Marquardt, A. Dahlmann, and P. Putthoff for excellent technical assistance.

